# PARP Inhibitor Counteracts Temozolomide Resistance in Glioblastoma Multiforme

**DOI:** 10.1101/2024.12.13.627927

**Authors:** Samarut Edouard, Tabourel Gaston, Briand Joséphine, Landais Yuna, Gratas Catherine, Loussouarn Delphine, Bougras-Cartron Gwenola, Cartron Pierre-François, Vallette François, Salaud Céline, Oliver Lisa, Aurelien A. Serandour

**Affiliations:** INSERM, CNRS, Nantes Université, Université d’Angers, CHU de Nantes, CRCI2NA, Nantes, France; Institut de Cancérologie de l’Ouest, 44805 Saint-Herblain, France; INSERM, Nantes Université, Ecole Centrale de Nantes, CR2TI, UMR 1064, Nantes, France

**Keywords:** Glioblastoma, Temozolomide, CRISPR screen, MMR, Olaparib

## Abstract

**Background:** Glioblastoma multiforme (GBM) is the most common malignant primary brain tumour in adults and is invariably associated with poor prognosis. Resistance to Temozolomide (TMZ), the standard chemotherapeutic agent, remains a major clinical challenge, particularly due to DNA mismatch repair (MMR) deficiencies. The aim of this study was to determine whether combining TMZ with the poly(ADP-ribose) polymerase inhibitor Olaparib (OLA) could overcome TMZ resistance in GBM.

**Methods:** We conducted *in vitro* experiments using U251 cell-line, including a TMZ-resistant derivative, and primary GBM cultures derived from patient tumours. A CRISPR/Cas9 knockout screen was employed to identify genes involved in TMZ resistance. Cell viability, proliferation, and morphology were assessed following treatment with TMZ, OLA, or their combination.

**Results:** The CRISPR screen identified inactivation of MMR pathway genes as key mediators of TMZ resistance. Co-treatment with OLA and TMZ demonstrated synergistic cytotoxicity in both parental and TMZ-resistant U251 cells, as well as in primary GBM cultures at diagnosis or relapse. Notably, OLA restored sensitivity to TMZ in MMR-deficient contexts and in tumours expressing O6-methylguanine-DNA-methyltransferase (MGMT). The combination treatment induced persistent DNA damage, cell cycle disruption, and cell death.

**Conclusions:** These findings provide strong preclinical evidence that combining TMZ with OLA can effectively overcome key mechanisms of TMZ resistance in GBM. This approach offers a promising therapeutic strategy warranting further clinical investigation.

**IMPORTANCE OF THE STUDY:** Temozolomide (TMZ) resistance remains a major therapeutic obstacle in glioblastoma (GBM), often driven by MMR deficiency or MGMT expression. While poly(ADP-ribose) polymerase (PARP) inhibitors have shown potential in other cancers, their role in overcoming TMZ resistance in GBM has remained unclear. In this study, a CRISPR screen identified MMR deficiency as a key driver of TMZ resistance. We further demonstrate that co-treatment with the PARP inhibitor Olaparib (OLA) restores TMZ sensitivity in both MMR-deficient and MGMT-expressing GBM cells and patient-derived cultures. These findings provide strong preclinical evidence supporting PARP inhibition as a promising therapeutic strategy to overcome chemoresistance in GBM and justify further clinical investigation.

**KEY POINTS:**

- PARP inhibitor Olaparib restores temozolomide sensitivity in resistant GBM cells
- Combination therapy overcomes resistance driven by MMR deficiency or MGMT expression
- Dual treatment induces persistent DNA damage and apoptosis in glioblastoma primary cultures

## INTRODUCTION

Glioblastoma, a WHO classification 4 grade tumour, is the most common and most aggressive malignant brain tumour in adults^1,2^. These highly proliferative gliomas mainly arise in the supratentorial brain^3^ and are characterized histologically by high cellularity, high mitotic activity, vascular proliferation, and necrosis^4^. GBM displays high inter and intra tumoral heterogeneity, which has been suggested to contribute to resistance to treatments ^5^. Indeed, tumour cells may differ in molecular, phenotypic, and functional characteristics, potentially enabling less sensitive tumour cell subpopulations to drive resistance to cancer drugs ^6^.

The standard of care for GBM involves surgical resection of the tumour if feasible, followed by a post-operative combination of concomitant radiotherapy and chemotherapy with Temozolomide (TMZ), known as the Stupp protocol^7^. Complete surgical resection is rarely achievable due to the GBM’s diffuse and infiltrative nature, as well as the risks of damaging surrounding brain tissue. This makes mandatory the use of complementary treatments^3^. TMZ is a monofunctional alkylating agent which induces cytotoxic lesions, including O^6^-methylguanine (O^6^MeG), N^7^-methylguanine (N^7^MeG) and N3-methyladenine (N3MeA) adducts ^8^. The DNA mismatch repair pathway (MMR), comprised of several proteins (MSH2, MSH3, MSH6, MLH1 and PMS2), is essential to mediate cytotoxicity of TMZ: mispairing of O6MeG with another thymine leads to repetitive rounds of MMR, known as “futile cycling”, ultimately inducing double-strand breaks and triggering p53-dependent cell cycle arrest and apoptosis^9,10^. These lesions are repaired by the O6-Methylguanine-DNA-Methyl Transferase (MGMT) enzyme, expressed by approximately 40% GBM tumours, leading to resistance to TMZ ^11–13^. However, in some GBM, the MGMT enzyme is silenced by promoter methylation, resulting in persistent O^6^MeG lesions and potential cytotoxicity. On the other hand, N^7^MeG and N^3^MeA adducts, which account for approximately 90% of methylation events induced by TMZ, are usually repaired by the base-excision repair (BER) pathway and contribute minimally to the overall cytotoxicity of TMZ^8,9^. The action of the BER pathway is coordinated by the Poly(ADP-Ribose) Polymerase (PARP) enzymes, which bind to the damaged DNA to recruit BER components^14,15^.

Despite aggressive treatment of GBM, including surgery and adjuvant radio-chemotherapy as discussed above, prognosis remains dismal with a median overall survival of 14-15 months and a five-year survival rate of only 5% ^16,17^. Almost all TMZ-sensitive GBM become drug resistant over time, resulting in tumour recurrence and clinical progression ^18^. Unfortunately, all the treatments used in tumour recurrence such as chemotherapy or targeted therapy failed to show significant clinical efficacy and survival improvement ^3^. Development of new therapeutic modalities therefore seems essential to improve care for patients with GBM.

One mechanism of acquired resistance to TMZ is the gain of mutations that inactivate the MMR pathway, thus inducing tolerance to DNA mispairing via O^6^MeG adducts, leading to cell survival ^19–21^. A way to overcome alkylating chemotherapy failure could be a targeting of the BER pathway, the inactivation of which could increase cytotoxicity of N^7^MeG and N^3^MeA adducts, otherwise repaired in normal cells and tumour cells with an intact BER pathway ^8,22^. The use of small molecule inhibitors of the PARP enzymes has shown promising results in preclinical models ^18,23–25^, through a potentiation of TMZ therapy by BER disrupting and other mechanisms such as destabilization of replication forks ^26^. Furthermore, it has been suggested that PARP inhibitors could also be effective in cancer cells expressing MGMT (without MGMT promoter methylation) ^27^, due to the necessary PARylation of MGMT by PARP for repairing TMZ-induced O^6^MeG adducts ^28^.

PARP1-inhibitors (PARP1i) have been marketed with specific indications with respect to homologous recombination repair (HR) status, which is considered a predictive biomarker in response to PARP1i^29^. In cells which have homologous repair deficiency (HRD), non-homologous end joining (NHEJ) would result from double strand break (DSB) formation, but with respect to homologous repair (HR), considered “error-free”, NHEJ affords DSBs repair with almost no consequence for sequence homology. The error-prone NHEJ promotes accumulation of DNA damage, and its activity is a major driver for PARP1i synthetic lethality in HR-defective cells^30^.

We present the results of *in vitro* experiments studying the use of the inhibitor of the enzyme Poly(ADP-ribose) polymerase 1 (PARP1), Olaparib (OLA), in combination with TMZ in the treatment of GBM tumour cells. Experiments were done on U251 cell lines, with no prior exposure to treatment, or on U251 resistant cell line due to prolonged exposure to TMZ. Then, experiments were replicated on several patient-derived cells at diagnosis or relapse.

## MATERIALS AND METHODS

### Materials

Temozolomide (LSC711) and Olaparib were obtained from Interchim (Montluçon, France), all other drugs used in this study were from Sigma (Saint Louis, MO). All cell culture products were obtained from Life Technologies (Carlsbad, CA).

### Cell culture

After informed consent, tumour samples classified as Glioblastoma type 4 (GBM) based on WHO criteria^31^, were obtained from patients undergoing surgical intervention at the “CHU de Nantes” and the “Tumorothèque IRCNA”. Within 4 h after surgical removal, GBM cells were recuperated after mechanical dissociation as described in *Brocard et al.*^32^. All procedures involving human participants were in accordance with the ethical standards of the ethic national research committee and with the 1964 Helsinki declaration and its later amendments or comparable ethical standards. GBM cells were cultured in defined medium (DMEM/F12 supplemented with 2 mM L-glutamine, N2 and B27 supplement, 2 µg/ml heparin, 20 ng/ml EGF, 40 ng/ml bFGF, 100 U/ml penicillin and 100 *µ*g/ml streptomycin). All the experiments with primary GBM cells were performed at early passages. Cells were regularly analysed for mycoplasma.

U251 cells were cultured in DMEM (Gibco, ref. 21969035) enriched with 10% FCS with 100 U/ml penicillin, 100 µg/ml streptomycin and 2 mM L-glutamine. U251 cell line was certified by Eurofins Genomics (Ebersberg, Germany). Cells were cultured at 37°C with 5% CO_2_ and 95% humidity. The cell medium was replaced every 2-3 days with fresh medium and the cells were reseeded in a new dish when the cell layer was approximately 80% confluent.

### CRISPR KO screen (cell culture)

We generated Cas9-expressing U251 clones by lentiviral transduction (Addgene, ref. 52962-LV) and blasticidine selection. We checked Cas9 expression by Western-Blot and selected the clone with the most efficient Cas9 nuclease activity using a Cas9 activity reporter (Addgene, ref. 67983 and ref. 67984). The Cas9-expressing U251 clone was expanded, transduced with the Brunello lentiviral library at MOI 0.3 (gift from David Root and John Doench; Addgene, ref. 73178-LV)^33^ and selected with puromycin. After expansion to 3 x 10^8^ cells: 10^8^ cells were harvested and frozen down, 10^8^ cells were treated with 50 µM TMZ and 10^8^ cells were treated with DMSO (vehicle condition). For the TMZ-treated cells, the medium was changed every 3 days with fresh TMZ. A minimum of 10^8^ DMSO-treated cells were replated every 3 days in fresh medium. The cells were cultured during 18 days, until the TMZ-treated cells reached the sub-confluence. At this point, DMSO-treated cells and TMZ-treated cells were harvested and frozen down.

### CRISPR KO screen (library preparation)

Genomic DNA was purified from cell pellets using Qiagen Blood & Cell Culture DNA Maxi Kit (ref. 13362). SgRNA sequences were amplified by PCR on genomic DNA using Q5 (NEB, ref. M0492S) and P5/P7 primers as recommended by Doench JG *et al.*^33^. Illumina libraries were quality-controlled by Agilent TapeStation, quantified by qPCR and submitted for HiSeq sequencing at the genomics core facility GenoA in Nantes, France.

### PARP1 inhibitor treatment

The PARP1 inhibitor, Olaparib (OLA) was dissolved in DMSO to give a stock concentration of 10 mM. In all experiments, the final working concentration of OLA was 1 µM (Supplementary Figure S8). The medium in the dishes was replaced every 2-3 days. DMSO was used as a vehicle control in all experiments (**Figure S1**).

### Transfection of U251 cells with siRNA

Cells were plated at 10^5^ cells/well in 6-well plates and 24h later transfected with 600 ng/ml siLuc, siMSH2, siMSH6 or siPMS2 (MISSION*^®^* esiRNA, Sigma-Aldrich, Saint-Louis, MI) using the using the Lipofectamine RNAiMax transfection reagent (Thermo-Fisher Scientific, Waltham, MA) for 48h and then used in experiments (**Figure S2**).

### Microscopy

Images of the cells were taken using an Axio Observer optical microscope (Zeiss®, Iéna, Germany) at 10x magnification. The images were then processed using ImageJ2 and Adobe Photoshop 8.0 software.

### Statistical analyses

Statistical analyzes were performed using one-way and two-way ANOVA tests with Prism 8 software. A p-value less than 0.05 is considered significant. The p-value of significant values is indicated by * < 0.05, ** < 0.01 and *** < 0.001.

## RESULTS

### Discovery of TMZ-resistance genes by CRISPR screen knock-out

Based on our observations from Rabé *et al*.^34,35^, we first conducted a CRISPR screen knock-out to identify genes involved in TMZ resistance. We generated a Cas9-expressing U251 clone and transduced it with the Brunello lentiviral library (gift from David Root and John Doench^33^). The cell library was expanded and treated with vehicle or with a therapeutic concentration of TMZ. About 95% of the cells died upon this TMZ insult. We harvested the cells once the surviving cells reached confluence after 18 days. After library preparation and sequencing of the sgRNA carried by the vehicle- and TMZ-treated cells, we analyzed the data using MAGeCKFlute ^36^ and found gene knock-outs significantly enriched in the TMZ-treatment condition, suggesting that these gene invalidations would confer resistance against TMZ. Several genes from the DNA MMR pathway were found: EXO1, MLH1, MSH2, MSH6, PMS1 and PMS2. We also found 3 other invalidations: BCL11A, CLHC1 and GPR75 involved respectively in neurone maturation, mitochondria and neurotransmission (**Figure 1A**). We did not find any genes whilst KO confers a higher sensitivity to the TMZ treatment.

**Figure 1:**
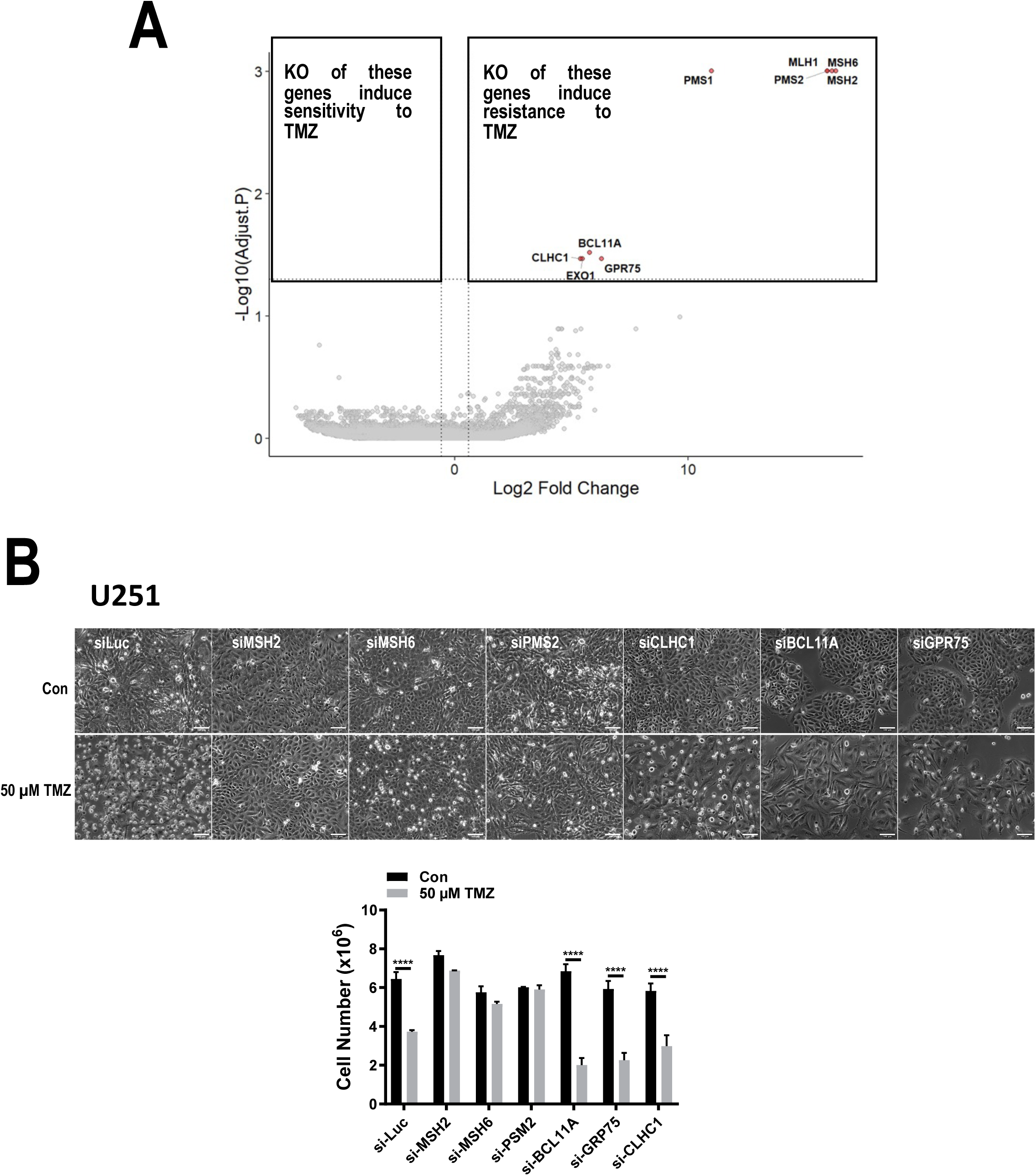
Genes involved in TMZ-resistance in U251 cells. **A.** CRISPR knock-out screen of U251 cells challenged with TMZ. **B.** SiRNA validation of genes identified by CRISPR screen. SiLuc = siLuciferase = control. Silencing of MSH2, MSH6 and PMS2 induce TMZ-resistance but not the silencing of CLHC1, BCL11A and GPR75.

We decided to proceed by validating the genes of interest by knock-down in U251 cells. We performed siRNA against the MMR members MSH2, MSH6, PMS2 and against BCL11A, CLHC1 and GPR75, using siRNA against Luciferase as control (**Figure 1B**). Only the knock-down of the MMR members MSH2, MSH6 and PMS2 conferred resistance against TMZ. This result suggests that, in the context of loss-of-function mutations, MMR deficiency is the main driver of TMZ resistance in GBM in recurrences.

### Combination of the PARPi OLA and TMZ counteracts TMZ-resistance in U251 cells

We tested the hypothesis that a PARPi could counteract TMZ resistance driven by MMR deficiency.

We found that 1 uM OLA had no effect on U251 cells in terms of cell proliferation, viability and morphology even after 16 days culture. In contrast, TMZ induced about a 90% cell death until day 12, after which the cells resumed proliferation as shown in Rabé *et al*.^34^, while the cells treated with OLA plus TMZ showed a marked decrease in cell number and there was no increase in cell number even after 16 days (**Figure 2A-B**).

**Figure 2:**
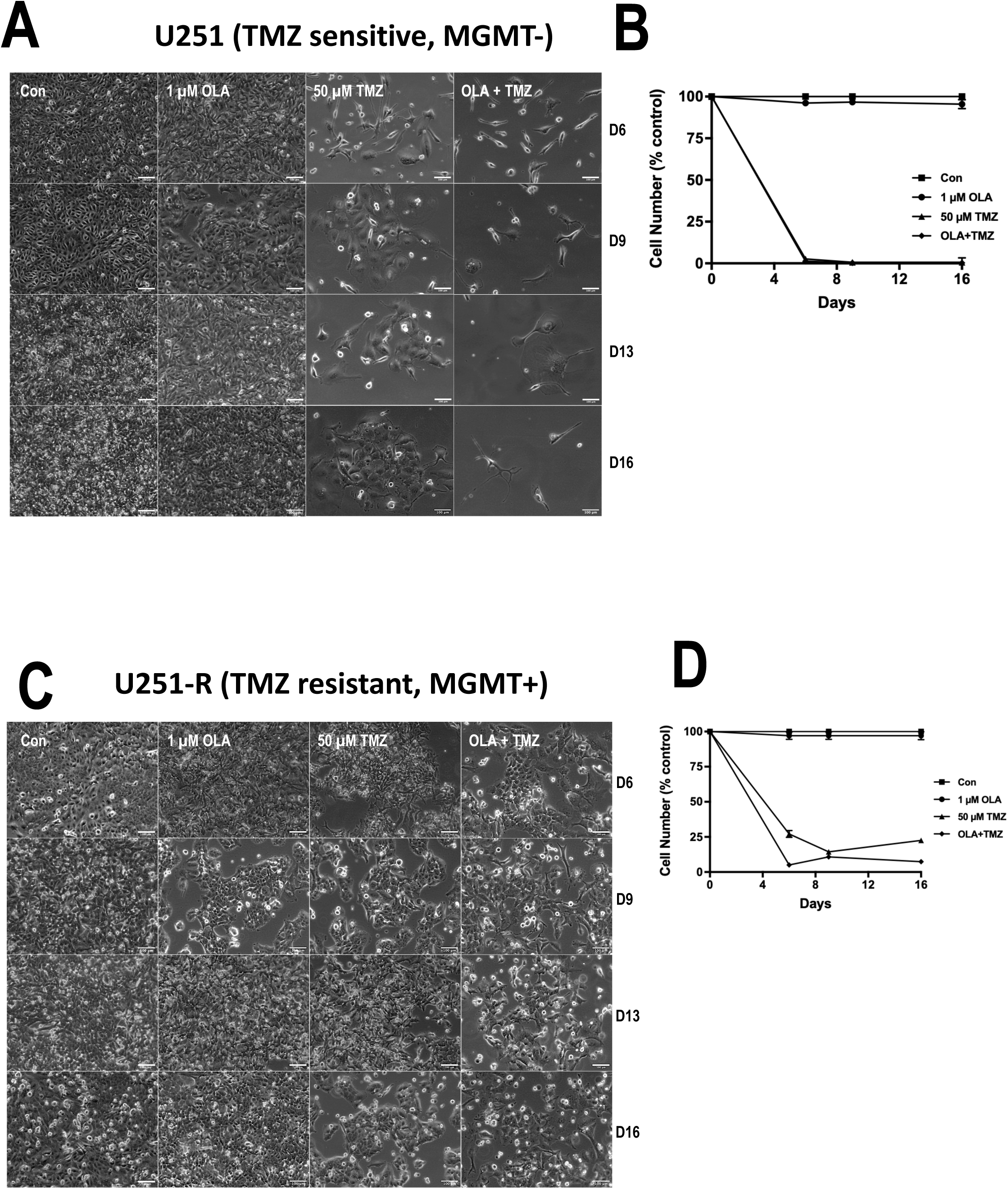
Effect of treatment of U251 and U251-R cells with Olaparib, TMZ or Olaparib + TMZ for 16 days.

We have shown that prolonged exposure to TMZ selects for resistant U251 by expressing MGMT *de novo* through promoter demethylation. These cells which we call U251-R cells were also treated with OLA and TMZ, and as shown in **Figure 2C-D**, we were able to induce greater cell death with OLA plus TMZ than with TMZ alone. OLA seems to have a synergistic effect when combined with TMZ.

We then questioned whether this was still true in a MMR-deficient context. We knocked-down MSH6 in U251 cells and treated the cells with control, OLA-only, TMZ-only or OLA plus TMZ (**Figure 3A-B**). We confirmed that MSH6 knock-down confers a strong resistance against TMZ. However, when combined to the TMZ, the OLA strongly reduces cell growth in MSH6-depleted cells. These results suggest that TMZ plus OLA have a synergistic effect that is able to counteract the TMZ-resistance induced by MSH6 deficiency.

**Figure 3:**
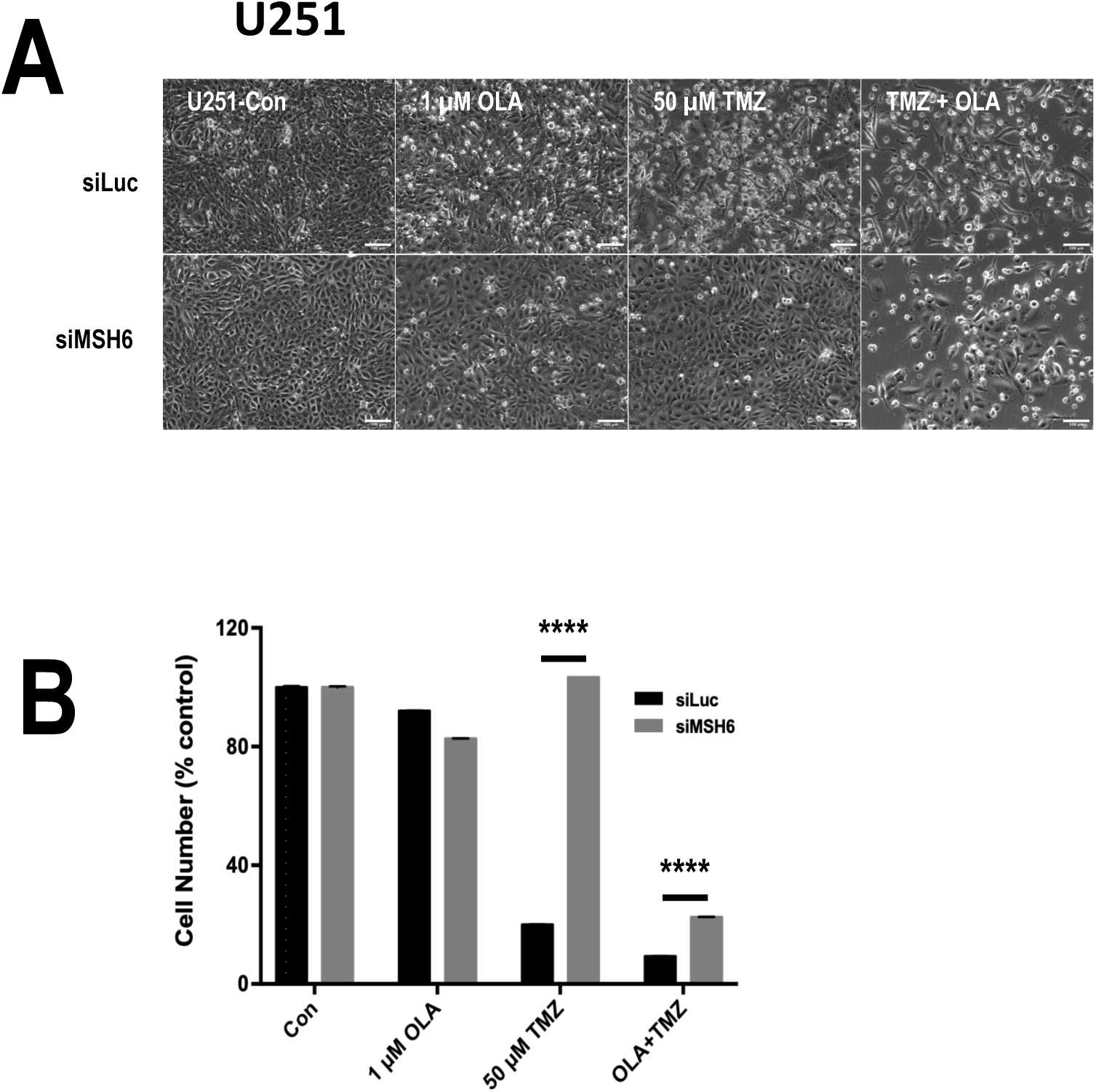
Effect of Olaparib treatment on U251 cells resisting TMZ. U251 cells were depleted in MSH6. This induces its resistance against TMZ. When Olaparib is added in combination with TMZ, cell death is restored.

### The combination of the PARPi OLA and TMZ counteracts TMZ-resistance in GBM primary cultures

We then tested the combination of TMZ plus OLA on glioblastoma primary cultures. We observed a synergistic effect of TMZ plus OLA in several patient cell cultures in GBM3, GBMG5, GBM69, GBM81, GBM120 (primary cultures at diagnosis), and GBMA1 and GBM96 (primary cultures at relapse) (**Figure 4A-B** and **Supplementary Figures S3-9**). In many cases after 17 days, the TMZ plus OLA treatment was able to eradicate all cells while the TMZ-only failed to do so. For the GBM86 patient, we cultured the GBM surgical excisions at diagnosis and at relapse, and treated them with TMZ-only, OLA-only or TMZ plus OLA (**Figure 4C-D**). Again, the combination of TMZ plus OLA had a stronger repressive effect on cell growth than TMZ alone both at diagnosis and relapse. It should be noted that the combination TMZ + OLA is also efficient in the context of MGMT expression like in U251-R, GBM3, GBMG5, GBM81, GBM96 and GBM120.

**Figure 4:**
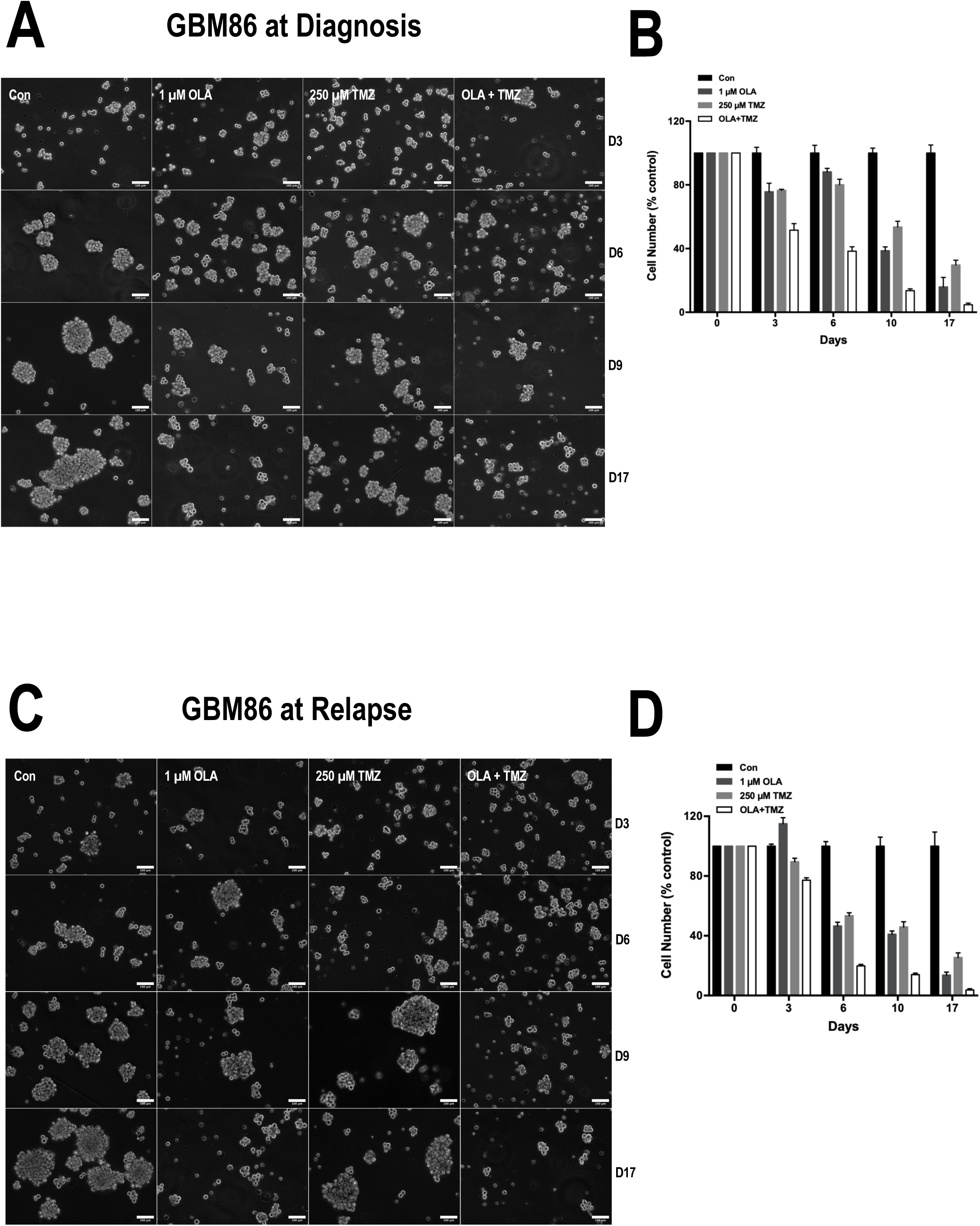
Effect of OLA, TMZ and OLA + TMZ on the GBM primary culture GBM86 at diagnosis and at relapse.

## DISCUSSION

Despite the use of TMZ as the standard chemotherapeutic agent for glioblastoma multiforme (GBM), resistance to this alkylating agent remains a major challenge, particularly at recurrence. In this study, we employed a genome-wide CRISPR/Cas9 knockout screen and identified loss of function in key mismatch repair (MMR) genes—*MSH2/6, MLH1*, *PMS1/2*—as the primary mechanism of resistance to TMZ. Notably, we show that co-treatment with the PARP inhibitor OLA can overcome this resistance not only in a TMZ-resistant GBM cell line but also in patient-derived GBM primary cultures. These findings unveil a therapeutically actionable vulnerability in TMZ-resistant GBM and support a new rationale for combining alkylating agents with PARP inhibitors in clinically relevant contexts.

Our CRISPR screen uncovered MMR gene disruption as the predominant hit conferring TMZ resistance. This finding corroborates prior work suggesting that functional MMR is essential for the cytotoxicity of TMZ-induced O^6^-methylguanine lesions^37^. During DNA replication, DNA polymerase mispairs O^6-methylguanine with thymine, prompting futile MMR cycles removing mismatched thymine and ultimately double-strand breaks and apoptosis^38^. Loss of MMR function abrogates this process, allowing cells to tolerate genotoxic insults. Indeed, Johnson *et al*.^39^ observed *MSH6* mutations in recurrent GBM following TMZ therapy, associating this with hypermutation and therapeutic failure. Our unbiased screen thus confirms in a systematic fashion that MMR loss is a major driver of TMZ resistance.

A key innovation of this study is the demonstration that OLA, a PARP1 inhibitor approved for clinical use in ovarian and breast cancers^40^, restores TMZ sensitivity in MMR-deficient cells. Previous studies have shown that PARP1 is essential for base excision repair (BER) and responds primarily to N7-methylguanine and N3-methyladenine lesions. These lesions comprise about 90% of TMZ-induced adducts but are usually non-cytotoxic^41^. Curtin^42^ proposed that blocking BER could render these adducts more deleterious, especially in cells that tolerate O^6MeG due to MGMT or MMR loss. Our results confirm this: OLA synergizes with TMZ and significantly reduces proliferation of MMR-deficient U251 cells, including those depleted of *MSH2, MSH6 or PMS2* as well as of a TMZ-resistant cells (U251-R). This synergy is consistent with prior findings by Higuchi et al. (2020)^43^, who showed that PARP inhibition could partially re-sensitize glioma cells lacking *MSH6* to TMZ. Moreover, our study extends these observations to patient-derived GBM cultures, including those from relapsed tumours with known therapy resistance. To our knowledge, this is the first demonstration of TMZ re-sensitization by a PARP inhibitor in *ex vivo* GBM cultures harbouring acquired resistance.

Our study reveals that PARP inhibition may represent a viable strategy to overcome MGMT-mediated resistance. Wu et al.^44^ recently reported that MGMT requires PARylation by PARP1 for its activity in repairing O^6MeG lesions. They showed that PARP inhibition impairs MGMT function, sensitizing otherwise resistant tumour cells. Erice et al.^45^ similarly found that melanoma cells with high MGMT expression were rendered TMZ-sensitive by PARP inhibition. In line with these reports, we show that OLA enhances TMZ efficacy in MGMT-expressing GBM models.

These findings have significant clinical implications. The majority of GBM patients with MGMT-unmethylated tumours do not benefit from TMZ therapy and face limited therapeutic options. Recent clinical trials of veliparib, a less potent PARP inhibitor, in combination with TMZ in newly diagnosed GBM failed to improve survival^46^. However, those studies did not stratify patients based on MMR status, nor did they include agents with stronger PARP trapping activity such as OLA or niraparib. Our data suggest that future trials should selectively enroll patients with MMR-deficient or MGMT-expressing tumours and employ more potent PARP inhibitors.

Additionally, there is growing interest in combining PARP inhibitors with radiotherapy. The PARADIGM-2 trial has explored OLA combined with radiation in newly diagnosed GBM and demonstrated good tolerability^47^. Combining OLA, TMZ, and radiotherapy could therefore represent a next-generation Stupp-like regimen tailored to MGMT-unmethylated or MMR-deficient GBM.

For patients with recurrent GBM, particularly those previously treated with TMZ, re-treatment is often ineffective due to resistance^48^. However, MMR mutations have been detected in such recurrences^37,39^, suggesting that our approach could benefit this large and underserved population. By showing efficacy in recurrent tumour cultures, we provide a strong preclinical rationale for evaluating OLA+TMZ in the recurrent setting.

The permeability of the Blood-Brain Barrier (BBB) remains a concern. Hanna *et al.*^49^ showed that OLA concentrations in brain tumours can vary, and optimizing delivery remains a challenge. Novel formulations or transient BBB disruption may be required to achieve therapeutic efficacy in patients. Finally, the hematologic toxicity of combining PARP inhibitors with TMZ must be carefully managed. Previous trials have shown manageable toxicity with adjusted dosing^50^, but further pharmacokinetic studies will be necessary to identify optimal dosing regimens that balance efficacy with tolerability.

## CONCLUSION

In summary, our study identifies MMR deficiency as the dominant mechanism of TMZ resistance in GBM via an unbiased CRISPR/Cas9 screen and demonstrates that PARP inhibition with OLA restores TMZ sensitivity in both MMR-deficient and MGMT-expressing tumours. These results redefine the landscape of TMZ resistance and suggest that a combination of TMZ and PARP inhibitors may have broad applicability in GBM therapy. Given the established clinical use of OLA and the limited options for resistant GBM, our findings justify immediate translational efforts and clinical evaluation of this strategy.

## ABBREVIATIONS LIST

BER: Base Excision Repair
GBM: Glioblastoma Multiforme
HR: Homology-directed Repair
MGMT: O^6^-Methylguanine-DNA-Methyl Transferase
N^3^MeA: N^3^-methyladenine
N^7^MeG: N^7^-methylguanine
NHEJ: Non-Homologous End Joining
O^6^MeG: O^6^-methylguanine
OLA: Olaparib
PARP: Poly(ADP-Ribose) Polymerase
TMZ: Temozolomide

## ACKNOWLEDGEMENTS

Funding: Ligue Contre le Cancer 44 ; Emergence Grant from Canceropole Grand Ouest.

## AUTHOR CONTRIBUTIONS

Samarut Edouard, Tabourel Gaston and Briand Joséphine contributed equally to the experimental work. Olivier Lisa and Sérandour Aurélien designed the project and wrote the manuscript with the involvement of Vallette François, Salaud Céline and Cartron Pierre-François. Landais Yuna, Gratas Catherine, Loussouarn Delphine and Bougras-Cartron Gwenola analysed the sequencing, histologic, clinical and qPCR data.

## DATA AVAILABILITY

The CRISPR screen sequencing data are available in ENA PRJEB73760. Other data will be made available upon reasonable request.

---

**Supplementary Figure S1:**
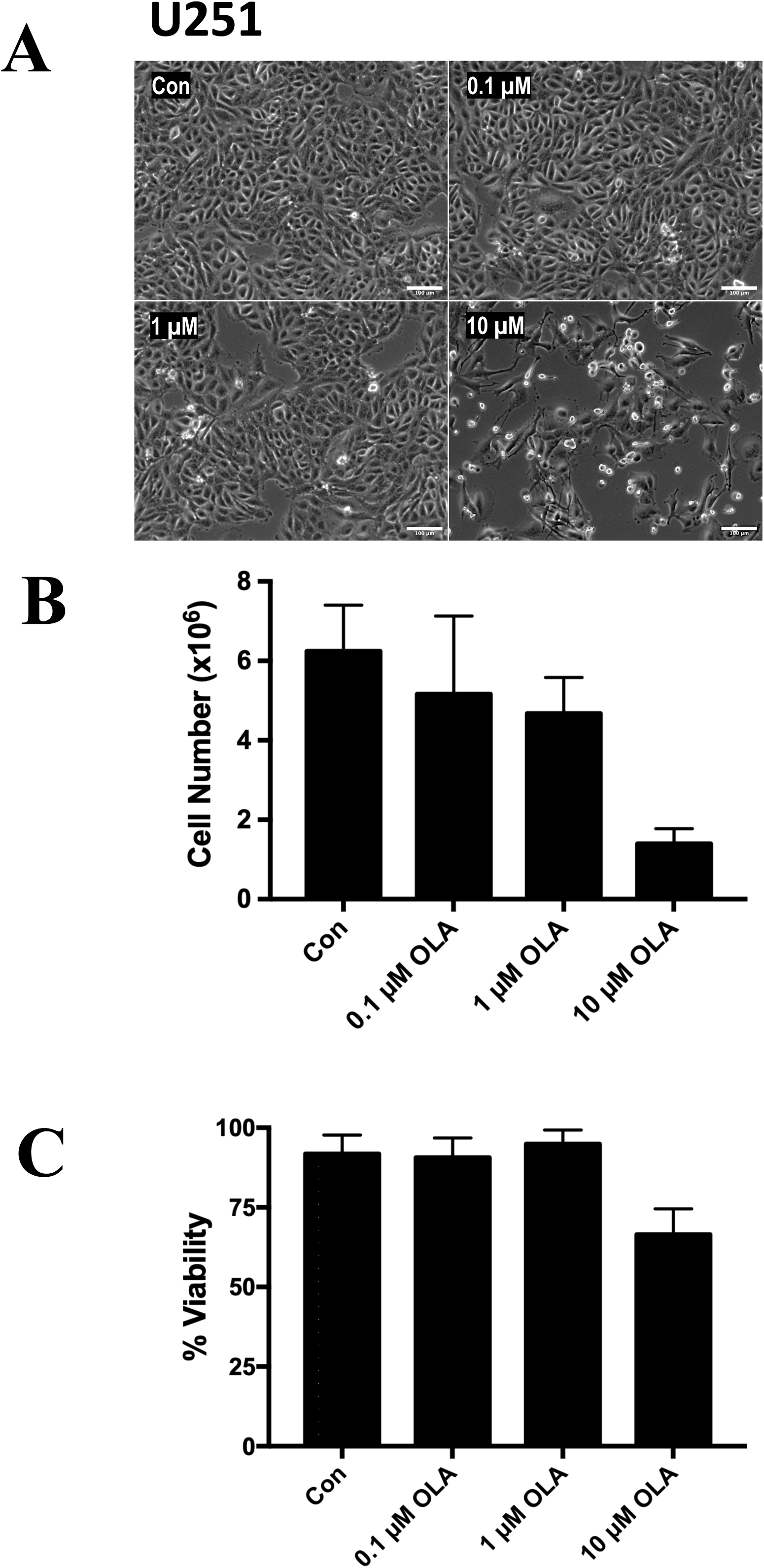
Effect of different OLA concentrations on U251 cells for 72 hours.

**Supplementary Figure S2:**
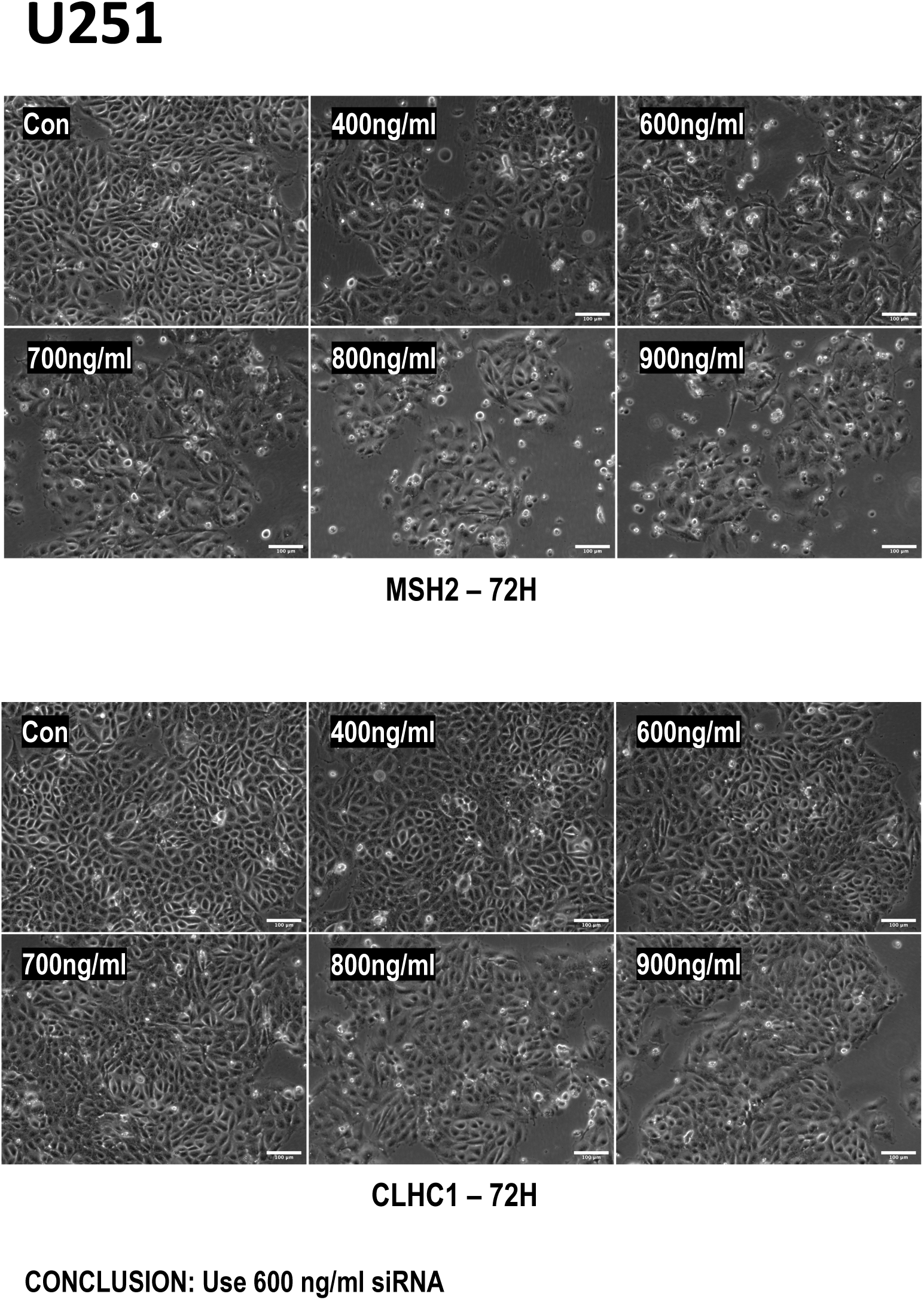
Effect of different siRNA concentration targeting MSH2 and CLHC1 on U251 cells for 72 hours.

**Supplementary Figure S3:**
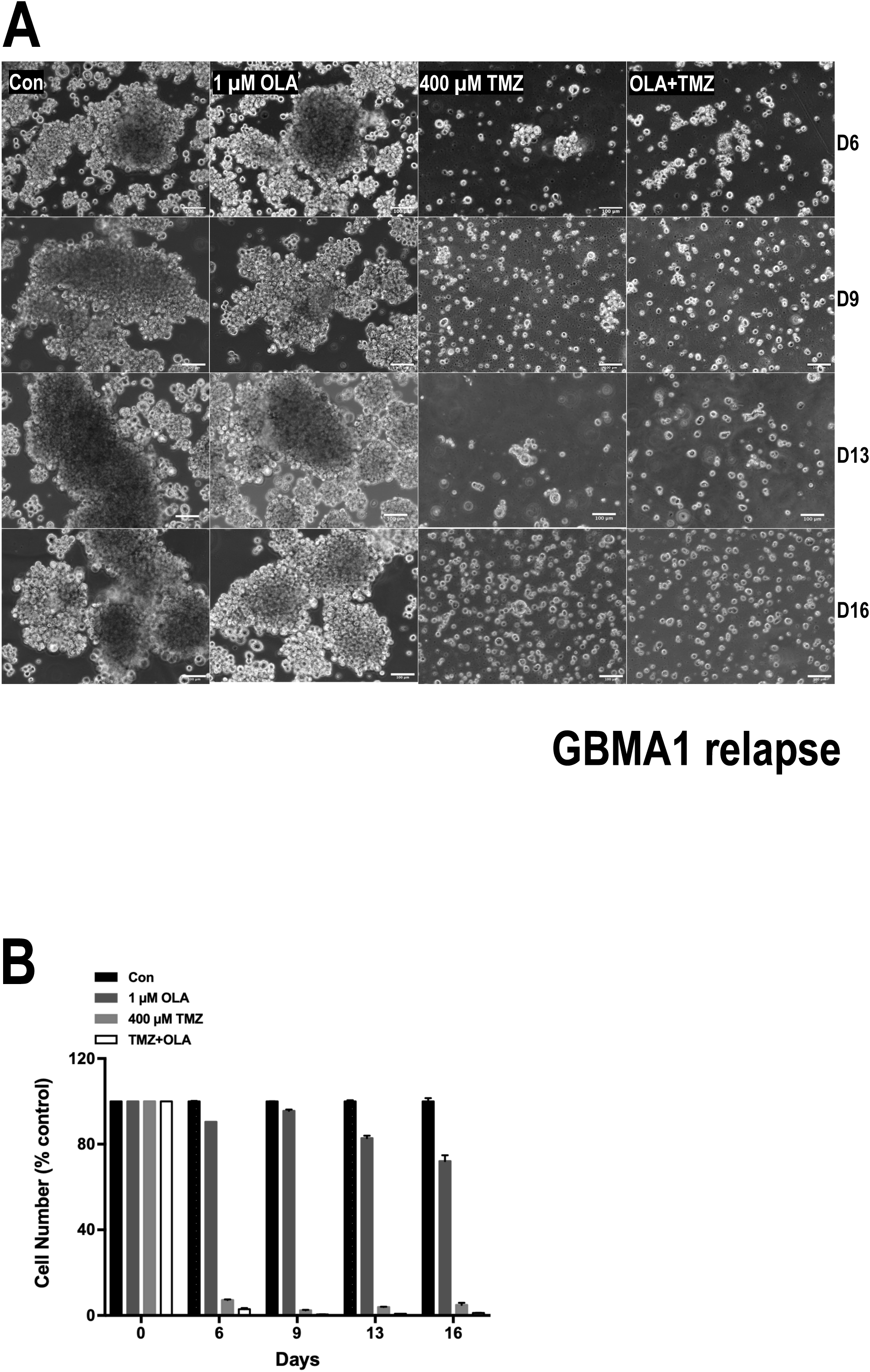
Effect of OLA, TMZ and OLA + TMZ on the GBM primary culture GBMA1 at relapse.

**Supplementary Figure S4:**
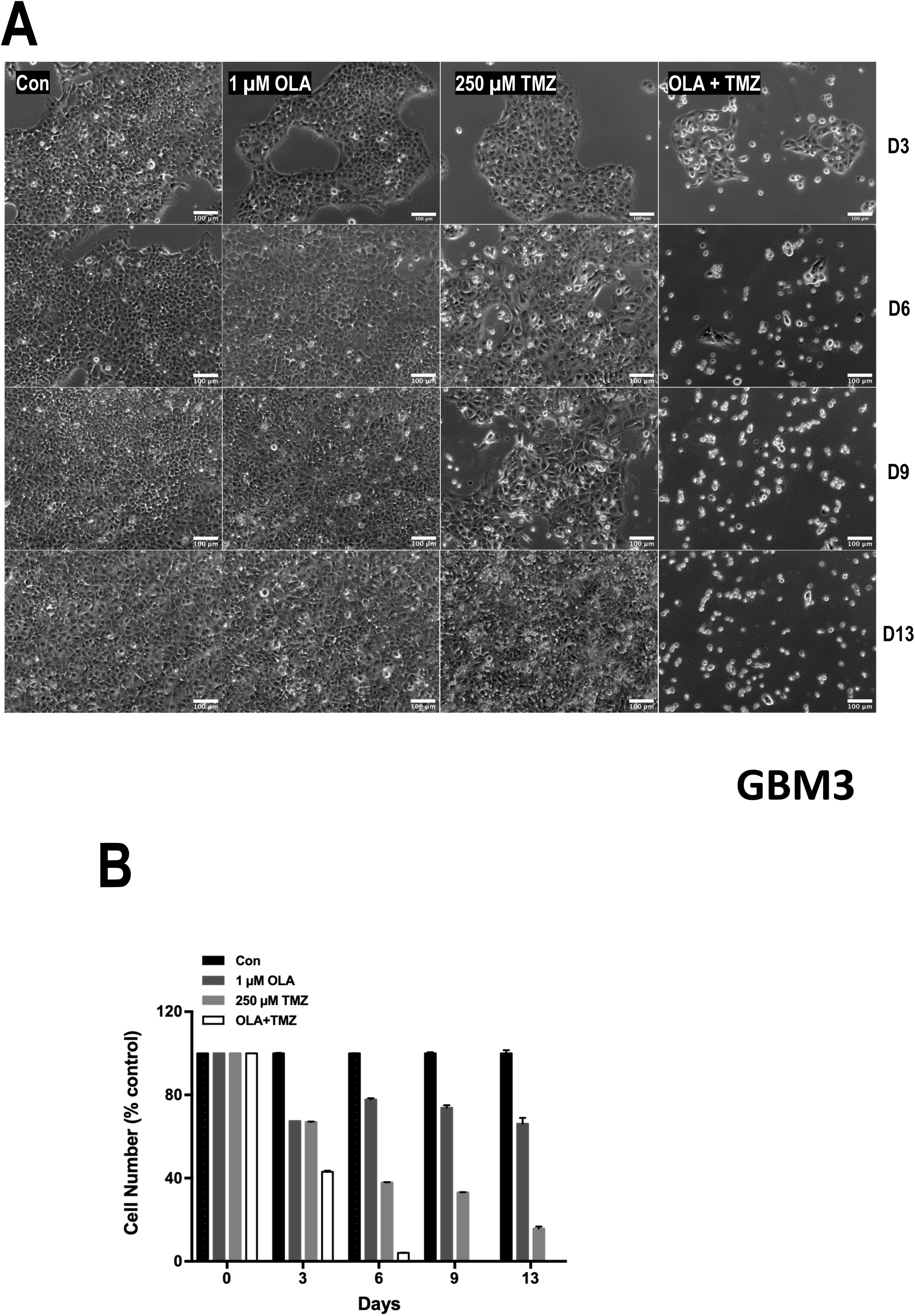
Effect of OLA, TMZ and OLA + TMZ on the GBM primary culture GBM3.

**Supplementary Figure S5:**
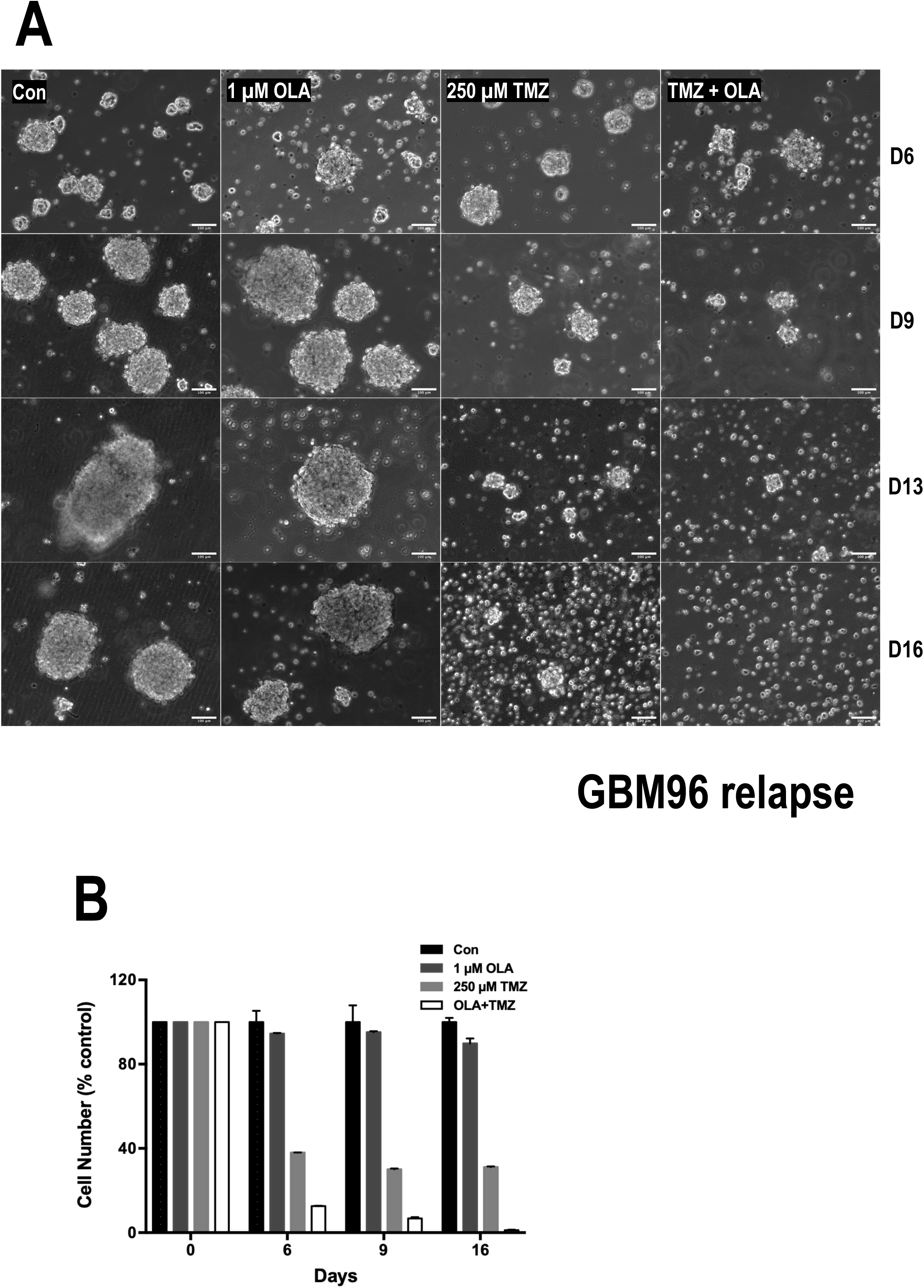
Effect of OLA, TMZ and OLA + TMZ on the GBM primary culture GBM96 at relapse.

**Supplementary Figure S6:**
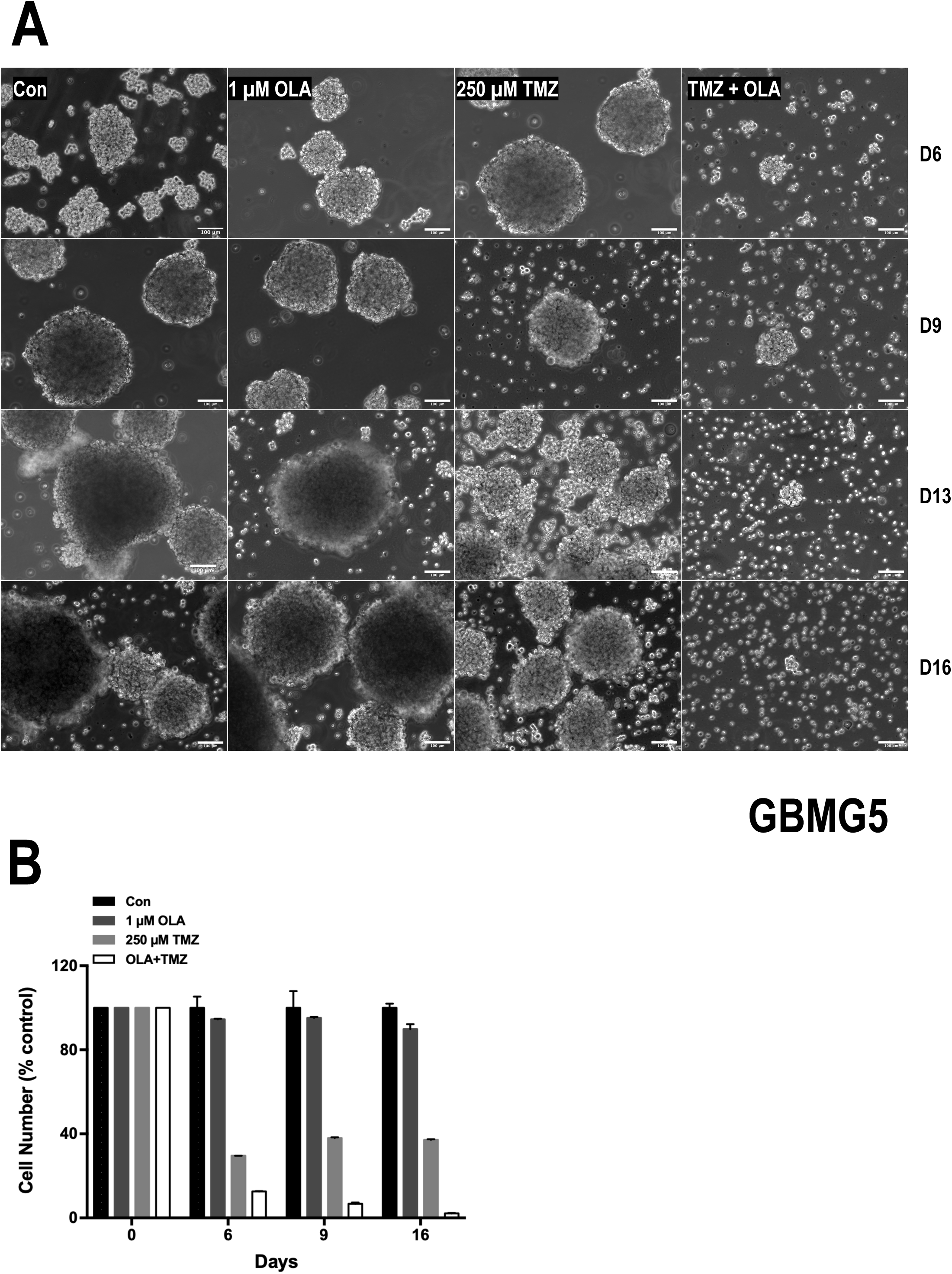
Effect of OLA, TMZ and OLA + TMZ on the GBM primary culture GBMG5.

**Supplementary Figure S7:**
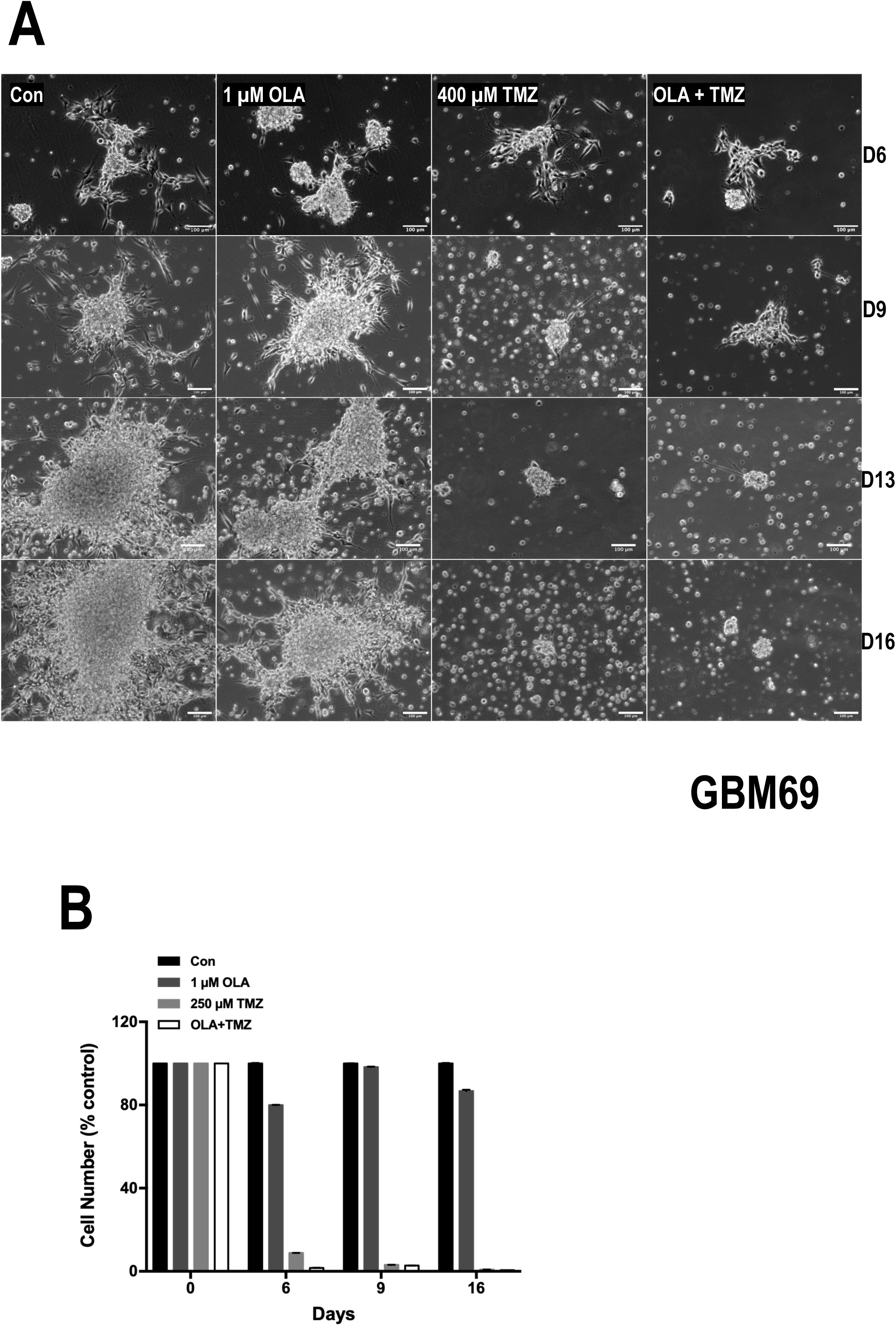
Effect of OLA, TMZ and OLA + TMZ on the GBM primary culture GBM69.

**Supplementary Figure S8:**
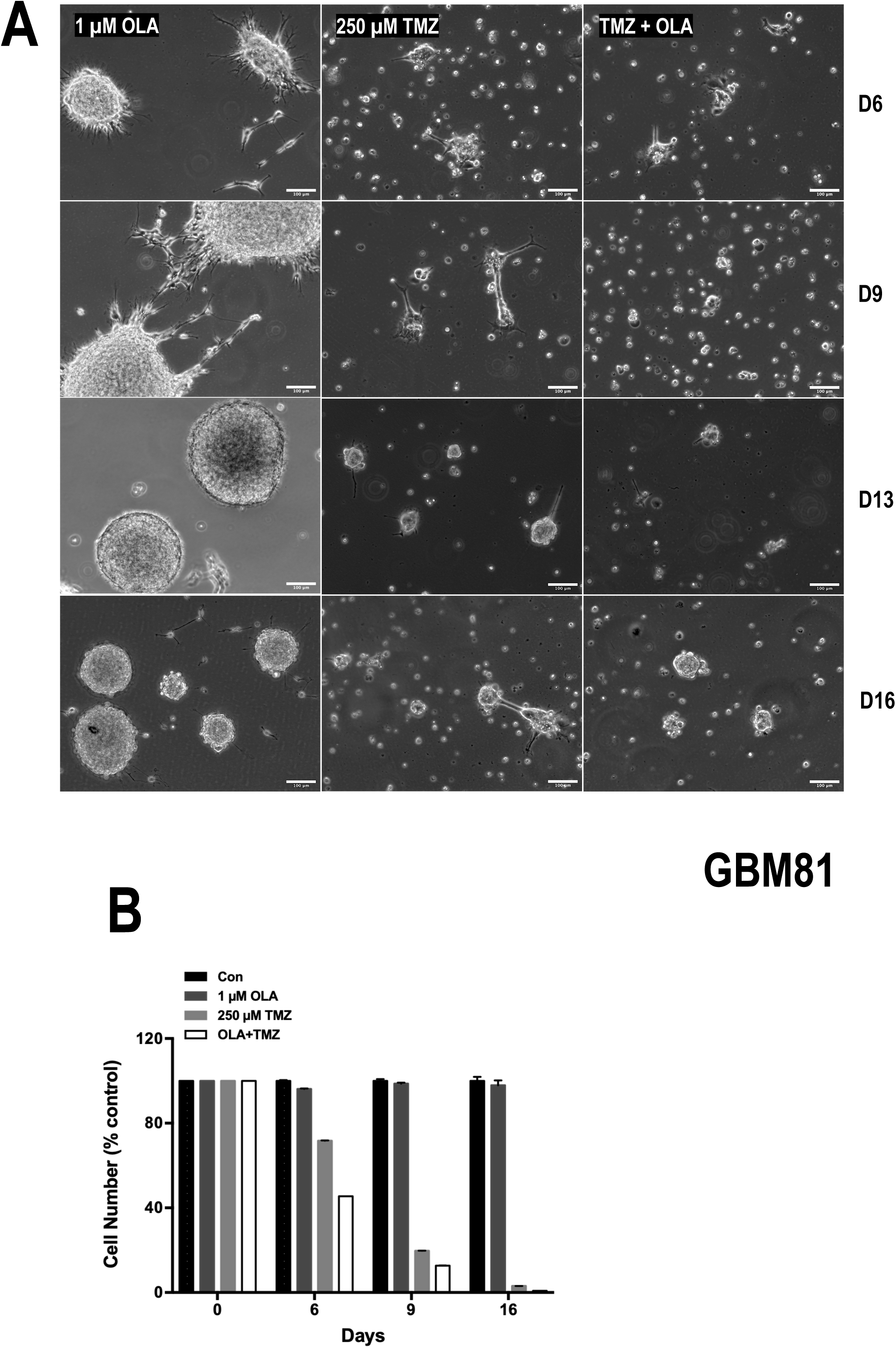
Effect of OLA, TMZ and OLA + TMZ on the GBM primary culture GBM81.

**Supplementary Figure S9:**
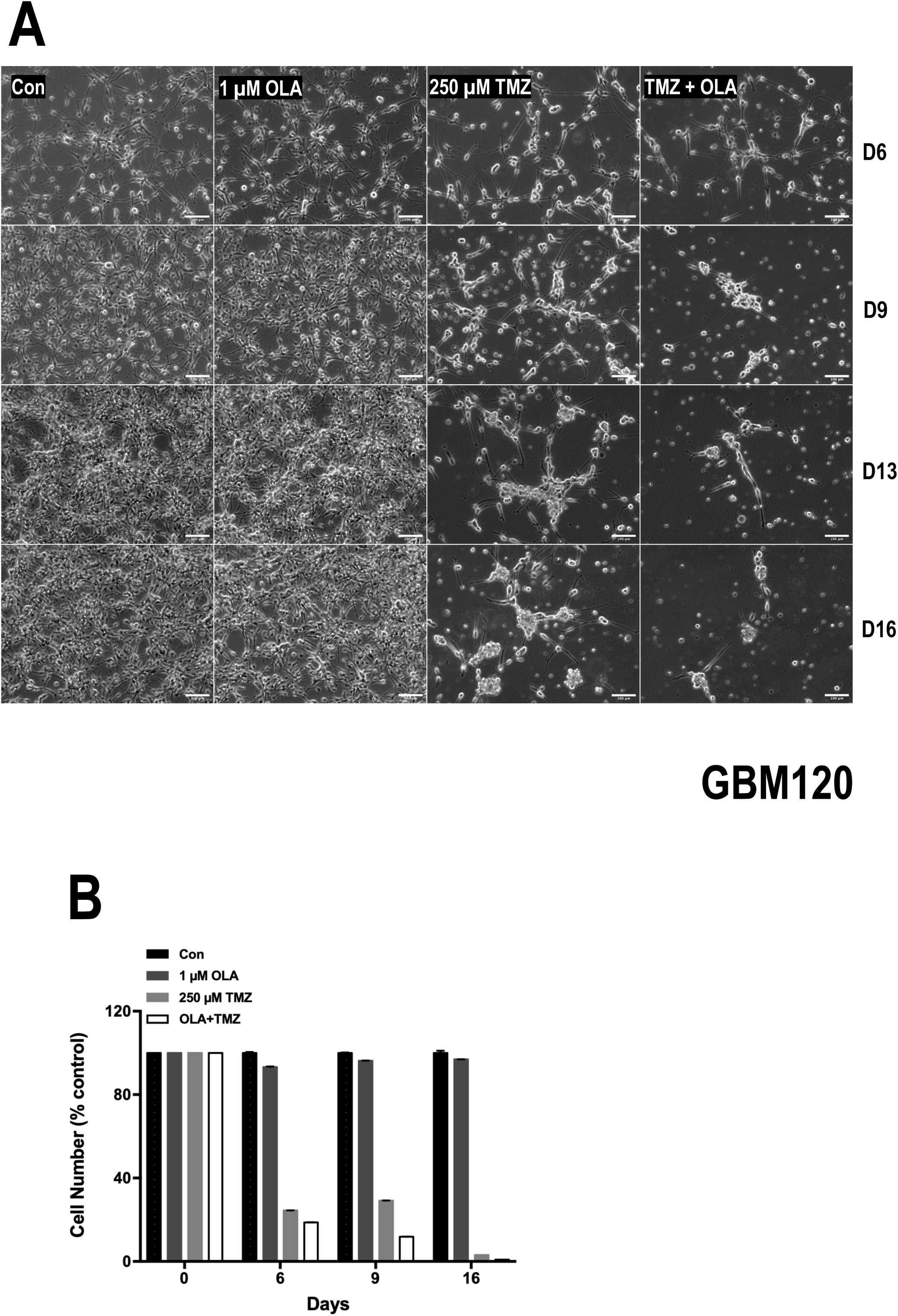
Effect of OLA, TMZ and OLA + TMZ on the GBM primary culture GBM120.

**Supplementary Table I:**
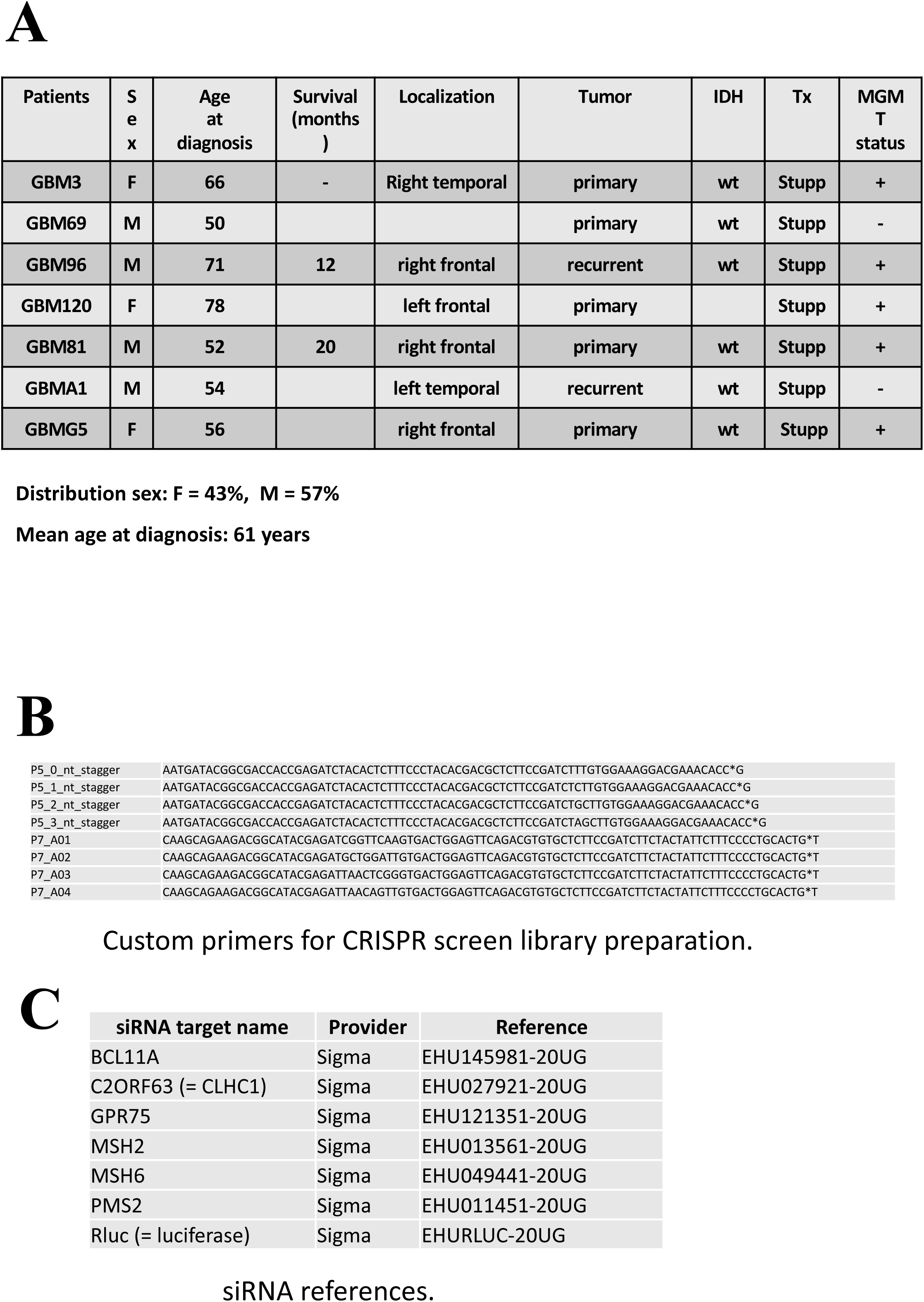
A. Table describing the GBM primary cultures used in this study B. Oligonucleotide sequences used to produce the CRISPR screen Illumina libraries C. Provider and references of the siRNA used.

## Notes

### Competing Interest Statement

The authors have declared no competing interest.

### Summary of Updates

The list of authors has been updated. The text has been deeply modified.

## REFERENCES

1 Louis DN, Perry A, Wesseling P, Brat DJ, Cree IA, Figarella-Branger D et al. The 2021 WHO Classification of Tumors of the Central Nervous System: a summary. Neuro Oncol 2021; 23: 1231–1251.

2 Thakkar JP, Dolecek TA, Horbinski C, Ostrom QT, Lightner DD, Barnholtz-Sloan JS et al. Epidemiologic and molecular prognostic review of glioblastoma. Cancer Epidemiol Biomarkers Prev 2014; 23: 1985–1996.

3 Oronsky B, Reid TR, Oronsky A, Sandhu N, Knox SJ. A Review of Newly Diagnosed Glioblastoma. Front Oncol 2021; 10. doi:10.3389/fonc.2020.574012.

4 Omuro A, DeAngelis LM. Glioblastoma and other malignant gliomas: a clinical review. JAMA 2013; 310: 1842–1850.

5 Inda M-M, Bonavia R, Seoane J. Glioblastoma Multiforme: A Look Inside Its Heterogeneous Nature. Cancers (Basel) 2014; 6: 226–239.

6 Cloughesy TF, Cavenee WK, Mischel PS. Glioblastoma: From Molecular Pathology to Targeted Treatment. Annual Review of Pathology: Mechanisms of Disease 2014; 9: 1–25.

7 Stupp R, Mason WP, van den Bent MJ, Weller M, Fisher B, Taphoorn MJB et al. Radiotherapy plus concomitant and adjuvant temozolomide for glioblastoma. N Engl J Med 2005; 352: 987–996.

8 Gupta SK, Smith EJ, Mladek AC, Tian S, Decker PA, Kizilbash SH et al. PARP Inhibitors for Sensitization of Alkylation Chemotherapy in Glioblastoma: Impact of Blood-Brain Barrier and Molecular Heterogeneity. Front Oncol 2018; 8: 670.

9 Sarkaria JN, Kitange GJ, James CD, Plummer R, Calvert H, Weller M et al. Mechanisms of Chemoresistance in Malignant Glioma. Clin Cancer Res 2008; 14: 2900–2908.

10 Hickman MJ, Samson LD. Role of DNA mismatch repair and p53 in signaling induction of apoptosis by alkylating agents. Proc Natl Acad Sci U S A 1999; 96: 10764–10769.

11 Hegi ME, Diserens A-C, Gorlia T, Hamou M-F, de Tribolet N, Weller M et al. MGMT gene silencing and benefit from temozolomide in glioblastoma. N Engl J Med 2005; 352: 997–1003.

12 Esteller M, Garcia-Foncillas J, Andion E, Goodman SN, Hidalgo OF, Vanaclocha V et al. Inactivation of the DNA-repair gene MGMT and the clinical response of gliomas to alkylating agents. N Engl J Med 2000; 343: 1350–1354.

13 Wick W, Weller M, van den Bent M, Sanson M, Weiler M, von Deimling A et al. MGMT testing--the challenges for biomarker-based glioma treatment. Nat Rev Neurol 2014; 10: 372–385.

14 Helleday T, Petermann E, Lundin C, Hodgson B, Sharma RA. DNA repair pathways as targets for cancer therapy. Nat Rev Cancer 2008; 8: 193–204.

15 Curtin NJ. PARP inhibitors for cancer therapy. Expert Rev Mol Med 2005; 7: 1–20.

16 Ostrom QT, Bauchet L, Davis FG, Deltour I, Fisher JL, Langer CE et al. The epidemiology of glioma in adults: a “state of the science” review. Neuro-Oncology 2014; 16: 896–913.

17 Stupp R, Hegi ME, Mason WP, van den Bent MJ, Taphoorn MJB, Janzer RC et al. Effects of radiotherapy with concomitant and adjuvant temozolomide versus radiotherapy alone on survival in glioblastoma in a randomised phase III study: 5-year analysis of the EORTC-NCIC trial. Lancet Oncol 2009; 10: 459–466.

18 Yuan AL, Meode M, Tan M, Maxwell L, Bering EA, Pedersen H et al. PARP inhibition suppresses the emergence of temozolomide resistance in a model system. J Neurooncol 2020; 148: 463–472.

19 Johnson BE, Mazor T, Hong C, Barnes M, Aihara K, McLean CY et al. Mutational analysis reveals the origin and therapy-driven evolution of recurrent glioma. Science 2014; 343: 189–193.

20 Wang J, Cazzato E, Ladewig E, Frattini V, Rosenbloom DIS, Zairis S et al. Clonal evolution of glioblastoma under therapy. Nat Genet 2016; 48: 768–776.

21 Yip S, Miao J, Cahill DP, Iafrate AJ, Aldape K, Nutt CL et al. MSH6 mutations arise in glioblastomas during temozolomide therapy and mediate temozolomide resistance. Clin Cancer Res 2009; 15: 4622–4629.

22 Bryant HE, Helleday T. Poly(ADP-ribose) polymerase inhibitors as potential chemotherapeutic agents. Biochem Soc Trans 2004; 32: 959–961.

23 Donawho CK, Luo Y, Luo Y, Penning TD, Bauch JL, Bouska JJ et al. ABT-888, an orally active poly(ADP-ribose) polymerase inhibitor that potentiates DNA-damaging agents in preclinical tumor models. Clin Cancer Res 2007; 13: 2728–2737.

24 Tentori L, Ricci-Vitiani L, Muzi A, Ciccarone F, Pelacchi F, Calabrese R et al. Pharmacological inhibition of poly(ADP-ribose) polymerase-1 modulates resistance of human glioblastoma stem cells to temozolomide. BMC Cancer 2014; 14: 151.

25 Hwang K, Lee J-H, Kim SH, Go K-O, Ji SY, Han JH et al. The Combination PARP Inhibitor Olaparib With Temozolomide in an Experimental Glioblastoma Model. In Vivo 2021; 35: 2015–2023.

26 Murai J, Huang SN, Das BB, Renaud A, Zhang Y, Doroshow JH et al. Differential trapping of PARP1 and PARP2 by clinical PARP inhibitors. Cancer Res 2012; 72: 5588–5599.

27 Erice O, Smith MP, White R, Goicoechea I, Barriuso J, Jones C et al. MGMT Expression Predicts PARP-Mediated Resistance to Temozolomide. Mol Cancer Ther 2015; 14: 1236–1246.

28 Wu S, Li X, Gao F, de Groot JF, Koul D, Yung WKA. PARP-mediated PARylation of MGMT is critical to promote repair of temozolomide-induced O6-methylguanine DNA damage in glioblastoma. Neuro-Oncology 2021. doi:10.1093/neuonc/noab003.

29 Rose M, Burgess JT, O’Byrne K, Richard DJ, Bolderson E. PARP Inhibitors: Clinical Relevance, Mechanisms of Action and Tumor Resistance. Front Cell Dev Biol 2020; 8: 564601.

30 Patel AG, Sarkaria JN, Kaufmann SH. Nonhomologous end joining drives poly(ADP-ribose) polymerase (PARP) inhibitor lethality in homologous recombination-deficient cells. Proc Natl Acad Sci USA 2011; 108: 3406–3411.

31 Louis DN, Perry A, Wesseling P, Brat DJ, Cree IA, Figarella-Branger D et al. The 2021 WHO Classification of Tumors of the Central Nervous System: a summary. Neuro Oncol 2021; 23: 1231–1251.

32 Brocard E, Oizel K, Lalier L, Pecqueur C, Paris F, Vallette FM et al. Radiation-induced PGE2 sustains human glioma cells growth and survival through EGF signaling. Oncotarget 2015; 6: 6840–6849.

33 Doench JG, Fusi N, Sullender M, Hegde M, Vaimberg EW, Donovan KF et al. Optimized sgRNA design to maximize activity and minimize off-target effects of CRISPR-Cas9. Nat Biotechnol 2016; 34: 184–191.

34 Rabé M, Dumont S, Álvarez-Arenas A, Janati H, Belmonte-Beitia J, Calvo GF et al. Identification of a transient state during the acquisition of temozolomide resistance in glioblastoma. Cell Death Dis 2020; 11: 19.

35 Rabé M, Fonteneau L, Oliver L, Morales-Molina A, Jubelin C, Garcia-Castro J et al. Cellular Heterogeneity and Cooperativity in Glioma Persister Cells Under Temozolomide Treatment. Front Cell Dev Biol 2022; 10: 835273.

36 Wang B, Wang M, Zhang W, Xiao T, Chen C-H, Wu A et al. Integrative analysis of pooled CRISPR genetic screens using MAGeCKFlute. Nat Protoc 2019; 14: 756–780.

37 Yip S, Miao J, Cahill DP, Iafrate AJ, Aldape K, Nutt CL et al. MSH6 mutations arise in glioblastomas during temozolomide therapy and mediate temozolomide resistance. Clin Cancer Res 2009; 15: 4622–4629.

38 Sarkaria JN, Kitange GJ, James CD, Plummer R, Calvert H, Weller M et al. Mechanisms of chemoresistance to alkylating agents in malignant glioma. Clin Cancer Res 2008; 14: 2900–2908.

39 Johnson BE, Mazor T, Hong C, Barnes M, Aihara K, McLean CY et al. Mutational analysis reveals the origin and therapy-driven evolution of recurrent glioma. Science 2014; 343: 189–193.

40 Lord CJ, Ashworth A. PARP inhibitors: Synthetic lethality in the clinic. Science 2017; 355: 1152–1158.

41 Gupta K, Burns TC. Radiation-Induced Alterations in the Recurrent Glioblastoma Microenvironment: Therapeutic Implications. Front Oncol 2018; 8: 503.

42 Curtin N. PARP inhibitors for anticancer therapy. Biochem Soc Trans 2014; 42: 82–88.

43 Higuchi F, Nagashima H, Ning J, Koerner MVA, Wakimoto H, Cahill DP. Restoration of Temozolomide Sensitivity by PARP Inhibitors in Mismatch Repair Deficient Glioblastoma is Independent of Base Excision Repair. Clin Cancer Res 2020; 26: 1690–1699.

44 Wu S, Li X, Gao F, de Groot JF, Koul D, Yung WKA. PARP-mediated PARylation of MGMT is critical to promote repair of temozolomide-induced O6-methylguanine DNA damage in glioblastoma. Neuro Oncol 2021; 23: 920–931.

45 Erice O, Smith MP, White R, Goicoechea I, Barriuso J, Jones C et al. MGMT Expression Predicts PARP-Mediated Resistance to Temozolomide. Mol Cancer Ther 2015; 14: 1236–1246.

46 Robins HI, Zhang P, Gilbert MR, Chakravarti A, de Groot JF, Grimm SA et al. A randomized phase I/II study of ABT-888 in combination with temozolomide in recurrent temozolomide resistant glioblastoma: an NRG oncology RTOG group study. J Neurooncol 2016; 126: 309–316.

47 Fulton B, Short SC, James A, Nowicki S, McBain C, Jefferies S, et al. PARADIGM-2: Two parallel phase I studies of olaparib and radiotherapy or olaparib and radiotherapy plus temozolomide in patients with newly diagnosed glioblastoma, with treatment stratified by MGMT status. Clin Transl Radiat Oncol 2018; 8: 12–16.

48 Stupp R, Mason WP, Van Den Bent MJ, Weller M, Fisher B, Taphoorn MJB et al. Radiotherapy plus Concomitant and Adjuvant Temozolomide for Glioblastoma. N Engl J Med 2005; 352: 987–996.

49 Hanna C, Kurian KM, Williams K, Watts C, Jackson A, Carruthers R et al. Pharmacokinetics, safety, and tolerability of olaparib and temozolomide for recurrent glioblastoma: results of the phase I OPARATIC trial. Neuro-Oncology 2020; 22: 1840–1850.

50 Kleinberg L, Supko JG, Mikkelsen T, Blakeley JO, Stevens G, Ye X et al. Phase I adult brain tumor consortium (ABTC) trial of ABT-888 (veliparib), temozolomide (TMZ), and radiotherapy (RT) for newly diagnosed glioblastoma multiforme (GBM) including pharmacokinetic (PK) data. JCO 2013; 31: 2065–2065.

